# Increased variability but intact integration during visual navigation in Autism Spectrum Disorder

**DOI:** 10.1101/2019.12.28.890004

**Authors:** Jean-Paul Noel, Kaushik J. Lakshminarasimhan, Hyeshin Park, Dora E. Angelaki

**Affiliations:** Center for Neural Science, New York University, New York City, NY, USA; Department of Neuroscience, Baylor College of Medicine, Houston TX, USA; Tandon School of Engineering, New York University, New York City, NY, USA

**Author notes:** Correspondence: Dr. Dora E. Angelaki, Center for Neural Science, Mayer 901, New York University, NY 10003.

## Abstract

Autism Spectrum Disorder (ASD) is a common neurodevelopmental disturbance afflicting a variety of functions from perception to cognition. The recent computational focus suggesting aberrant Bayesian inference in ASD has yielded promising but conflicting results in attempting to explain a wide variety of phenotypes by canonical computations. Here we used a naturalistic visual path integration task that combines continuous action with active sensing and allows tracking of subjects’ dynamic belief states. Both groups showed a previously documented bias pattern, by overshooting the radial distance and angular eccentricity of targets. For both control and ASD groups, these errors were driven by misestimated velocity signals due to a non-uniform speed prior, rather than imperfect integration. We tracked participant’s beliefs and found no difference in the speed prior, but heightened variability in the ASD group. Both end-point variance and trajectory irregularities correlated with ASD symptom severity. With feedback, variance was reduced and ASD performance approached that of controls. These findings highlight the need for both more naturalistic tasks and a broader computational perspective to understand the ASD phenotype and pathology.

## Introduction

Autism Spectrum Disorder (ASD) is a heterogeneous neurodevelopmental disorder with high prevalence (Xu et al., 2018). In response to its pervasiveness, researchers have recently turned their attention to computational and normative tools attempting to identify canonical computations underlying ASD symptomatology (e.g., Robertson & Baron-Cohen, 2018; Rosenberg et al., 2015). A promising candidate family is that of probabilistic inference (Doya et al., 2007), and indeed a large number of Bayesian accounts of ASD have recently been put forward – positing an anomaly in the strength of Bayesian priors (Pellicano and Burr, 2012; Friston et al., 2013), the abnormal updating of these priors (Lawson et al., 2017; Lieder et al., 2019), the aberrant precision in sensory representations (Brock, 2012; Lawson et al., 2015; Zaidel et al., 2015; Karvelis et al., 2018), and the atypical weighting of sensory prediction error (Friston et al., 2013; Haker et al., 2016; van de Cruys et al., 2016).

Unfortunately, as for many other aspects of the ASD phenotype and pathology, there is yet no clear consensus. Remarkably, both attenuated (Karaminis et al., 2016; Noel et al., 2016) and intact (Pell et al., 2016; Croydon et al., 2017; Karvelis et al., 2018) priors have been reported, as well as normal (Manning et al., 2017) and abnormal (Lawson et al., 2017; Lieder et al., 2019) updating of these priors. These conflicting results may partially be due to a lack of quantification. Many studies have based their conclusions on a loose link to “reduced top-down modulation of sensory processing’’, “difficulties in accessing underlying statistical rules in an unstable context” or “impairment in predictive abilities” without any quantitative fit of the inference/predictive process (Croydon et al., 2017; Gonzalez-Gadea et al., 2015; Noel et al., 2016; Palmer et al. 2015; Pell et al., 2006; Robic et al., 2015; Sinha et al., 2014; Skewes et al., 2015; Skewes & Gebauer, 2016; Turi et al., 2015, 2016). A handful of studies have provided Bayesian model simulations (Karaminis et al., 2016; Powell et al., 2016; Zaidel et al., 2015). To our knowledge, the only study that has computationally disentangled individuals’ likelihoods and priors (Karvelis et al., 2018) reported no difference in prior distributions between ASD and control groups.

In addition to a lack of quantification, another contributing factor to conflicting conclusions may be the widespread use of constrained and data-poor tasks defined by binary behavioral outcomes. Separating perception from action, for example, is a laboratory construct that has little to do with the challenges of everyday experiences. Furthermore, binary outcome tasks offer few data points to allow firm exploitation of complex computations, like fitting likelihoods and priors. We argue that, to understand the dynamic neural processes that mediate natural behavior – and deficits thereof - we must study recurrent neural computations by using continuous-time behavioral outcomes where actions influence sensory inflow — particularly when crucial variables cannot be directly observed such that the observer must draw inferences about those latent variables, as is often the case in ecological behaviors.

Here we used a virtual reality navigation task that allows exploitation of brain computation in the naturalistic setting of continuous action and active sensing, as well as dynamic on-line inference about latent, task-relevant variables (Lakshminarasimhan et al., 2018). More specifically, we use a virtual navigation task that required control and ASD participants to use a joystick to actively acquire memorized targets by integrating visual motion cues (optic flow). This task is not only more natural in terms of the dynamic, closed-loop interactions between sensory inflow, internal beliefs, and actions, but also requires the continuous integration of visual motion cues – a process previously reported to be abnormal in ASD (Spencer et al., 2000; Milne et al., 2002; Pellicano et al., 2005), but recently suggested to reflect heightened sensitivity to noise (Zaidel et al., 2015). Further, our navigate-to-target task provides a rich and continuous dataset (i.e., two-dimensional movement trajectories extending for ∼5 seconds/trial) permitting the tracking of belief states (Lee et al., 2014; Lakshminarasimhan et al., 2018), and efficient fitting of different components forming Bayesian computations (i.e., priors and likelihood functions).

## Results

We asked ASD (n = 14) and matched control (n = 25, see *Methods* for detail) adolescents to use a joystick to virtually navigate toward and stop at the location of a briefly presented visual cue, a “firefly”. No landmarks were presented, only ground-plane triangular elements providing optic flow cues (**Figure 1A**). In the second half of trials participants were instructed in the task via feedback in the form of concentric circles indicating the location of the target, and a colored arrow (green if “rewarded” and red if “unrewarded”, **Figure 1B**). The portion of space “rewarded” was adaptively manipulated to become more restrictive with improved task performance (see *Methods*). Targets were distributed randomly and uniformly (**Figure 1C**) within a range of r = 1-6 meters (r, radial distance) and θ = ± 42.5° (θ, angular eccentricity) of visual angle relative to where the subject was stationed at the beginning of the trial. This radial distance is within the regime where humans are known to overshoot targets (Lakshminarasimhan et al., 2018; undershooting appears at and beyond approximately 20m). Indeed, visualizing exemplar target locations and trial trajectories (**Figure 1C**) during the block without feedback suggests that participants explored a larger space than that required by target locations. To further depict this trend, we expressed participants’ responses in polar coordinates (**Figure 1D**), with an eccentricity from vertical (angular response, 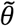) and a radial distance 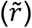. In the example presented in **Figure 1D** (left), the error vector points radially outward and away from straight ahead. This pattern was consistent across trials for this particular subject, as shown in the vector field of errors (**Figure 1D**, right). This profile of errors implies consistent overshooting both in terms of absolute distance traveled and angular rotation. We initially focus on performance and impact of feedback on path integration for control subjects. Then, we assess baseline performance and the impact of feedback in individuals with ASD, as compared to neurotypical individuals.

**Figure 1.**
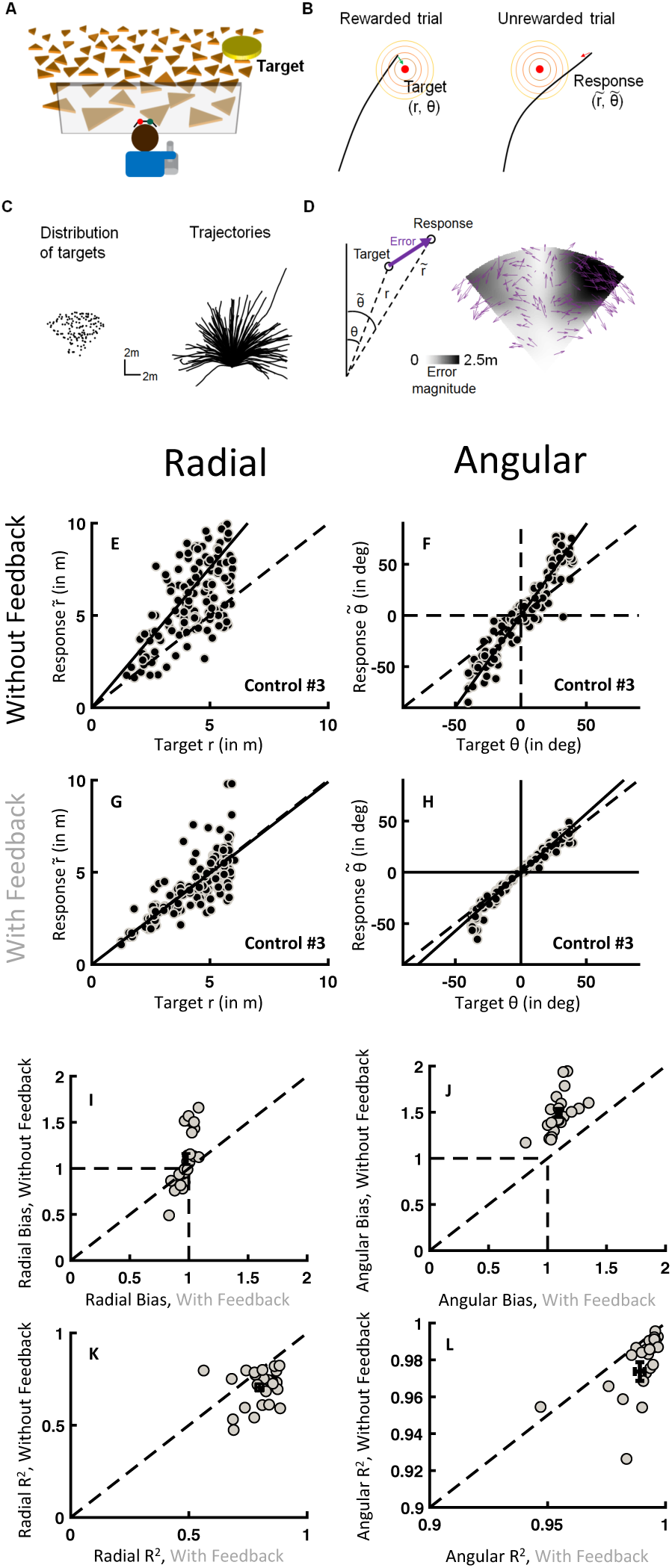
Experimental Protocol and Normal Performance. **A)** Participants use a joystick to navigate to a flashed target (yellow disc, or “firefly”) using optic flow generated by ground-plane triangles. **B)** Example trajectory of a participant approaching the unseen target. In the feedback block, after participants have made their response, concentric circles and either a green (if “rewarded”) or red (if “unrewarded”) arrow appears indicating the true location of the target. **C)** Distribution of targets (left) and example trajectories from one experimental block (right). **D)** Left: Target and endpoints expressed in polar coordinates (angular distance: θ; radial distance: r). Right: Errors of the example trajectories. **E-H)** Scatter plots of radial and angular distance responses (y-axis) as a function of the respective target distance (x-axis) for a representative subject (Control Subject #3), shown separately without and with feedback. Individual dots are single trials. Solid lines: linear regression; Dashed lines: identity lines. **I-L)** Scatter plots of regression slopes (1 = no bias, <1 = undershooting, >1 = overshooting) for all participants individually (gray dots) and population average (error bars: ± SEM).

### Typical Performance and Impact of Feedback

To quantitatively assess the apparent under-estimation in self-position (and thus overshooting) during path integration, we separately compared the radial and angular error by performing a linear regression between target positions (*r, θ*) and responses 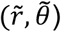. **Figures 1E** and **F** show these regressions, respectively in the radial and angular dimension, for an exemplar control individual during the block without feedback. The linear fits account relatively well for the pattern of responses observed (Radial: R^2^ = 0.54; Angular: R^2^ = 0.92), while also evidencing considerable variability across trials, particularly in the radial dimension (**Figure 1E**, individual dots are single trials). Further, these data suggest that bias during path integration is multiplicative: the greater the distance traveled, the greater the error, as indicated by regression slopes over 1 (slope = 1 reflects no bias. Radial: 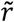 vs. *r* slope = 1.51, Angular: 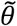 vs. *θ* slope = 1.78). There is no notable distance-independent bias, which would have been expressed as regressions with non-zero intercepts. **Figures 1G** and **H** illustrate the relationship between target location and responses in the block with feedback (same participant as in **Figures 1E-F**), showing dramatic improvement in terms of both accuracy (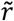 vs. *r* slope = 0.99, 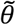 vs. *θ* slope = 1.14) and precision of endpoints (Radial: R^2^ = 0.77; Angular: R^2^ = 0.98) compared to the block without feedback.

The above findings were consistent across subjects, as illustrated in **Figures 1I** and **J**. In the absence of feedback, both radial (mean ± standard error of the mean (SEM): 1.10 ± 0.06, p = 0.045) and angular (1.49 ± 0.05, p = 1.50 × 10^−14^) overshooting biases were appreciable. Similar to Lakshminarasimhan et al., 2018, the effect size was 4 to 5 times greater in the angular (50%) than radial (10%) dimension (78% vs. 19% in Lakshminarasimhan et al., 2018): all subjects show angular over-estimation, while not all show radial over-estimation.

The introduction of feedback completely eliminated overshooting in radial distance (0.97 ± 0.02, p = 0.6, **Figure 1I**) and reduced, but not completely eliminated, the angular bias (1.09 ± 0.02, p = 5.51 × 10^−8^; **Figure 1J**). Interestingly, providing feedback at the end of each trial increased not only accuracy, but also precision (feedback – without feedback, ΔR^2^ Radial = 0.09 ± 0.02, p = 4.25 × 10^−4^; ΔR^2^ Angular = 0.01 ± 0.004, p = 0.003: **Figure 1K** and **L**). Further, the introduction of feedback drove participants to faster trajectories, both in the radial (F = 125.0, p<0.001) and angular (F=24.0, p<0.001) dimension, and in turn to shorter trials (from 5.42 ± 0.05 without feedback, to 3.70 ± 0.05 seconds with feedback, F=98.22, p<0.001; **Figure S1**).

### Path Integration Improves with feedback due to a reduction in self-motion uncertainty

To further understand the root cause of participant’s overshooting of the target, and most importantly, the driving, latent, mechanism behind their improvement during trial-to-trial feedback, we instantiated two dynamic Bayesian observer models (first introduced in Lakshminarasimhan et al., 2018). Both models assume that subjects maintain estimates of both the mean and uncertainty associated with their location, and steer toward the target to maximize reward on each trial (see Lakshminarasimhan et al., 2018 and *Methods*). The trajectories generated by each model correspond to the subject’s beliefs about their distance to target throughout the trial. Thus, these models are fitted to the whole movement trajectory for each trial by maximizing the overlap between the posterior distribution over believed position and the target region at the end of each trial (see *Methods*).

The first model hypothesizes that participants overshoot targets because they misestimate their speed due to a non-uniform prior that biases velocity estimates. In contrast, sensory evidence accumulation (path integration) is assumed to be lossless (**Figure S2A;** Hurlimann et al., 2002; Stocker & Simoncelli, 2006; Weiss et al., 2002; Petzschner & Glasauer, 2011). The inference of velocity estimates from optic flow signals depends on the shape of the observer’s prior distribution for velocity (**Figure 2A**, solid black line, *a*_*v*_ and *a*_*w*_ defining the exponential decay of the radial and angular velocity components, respectively), which is combined with a likelihood function (**Figure 2A**, dashed black line, *b*_*v*_ and *b*_*w*_, respectively, defining the linear relationship between velocity and the variance of the radial and angular likelihood, see *Methods*) to generate the final posterior estimate of velocity (**Figure 2A**, green). Unlike Lakshminarasimhan et al., 2018, here the exponents dictating the shape of the prior distribution (*a*_*v*_ and *a*_*w*_) were allowed to be negative or positive (allowing for both under- and over-estimation), given that participants are overall accurate during the feedback condition, and even in the block without feedback a subset of participants did not radially overshoot targets.

**Figure 2.**
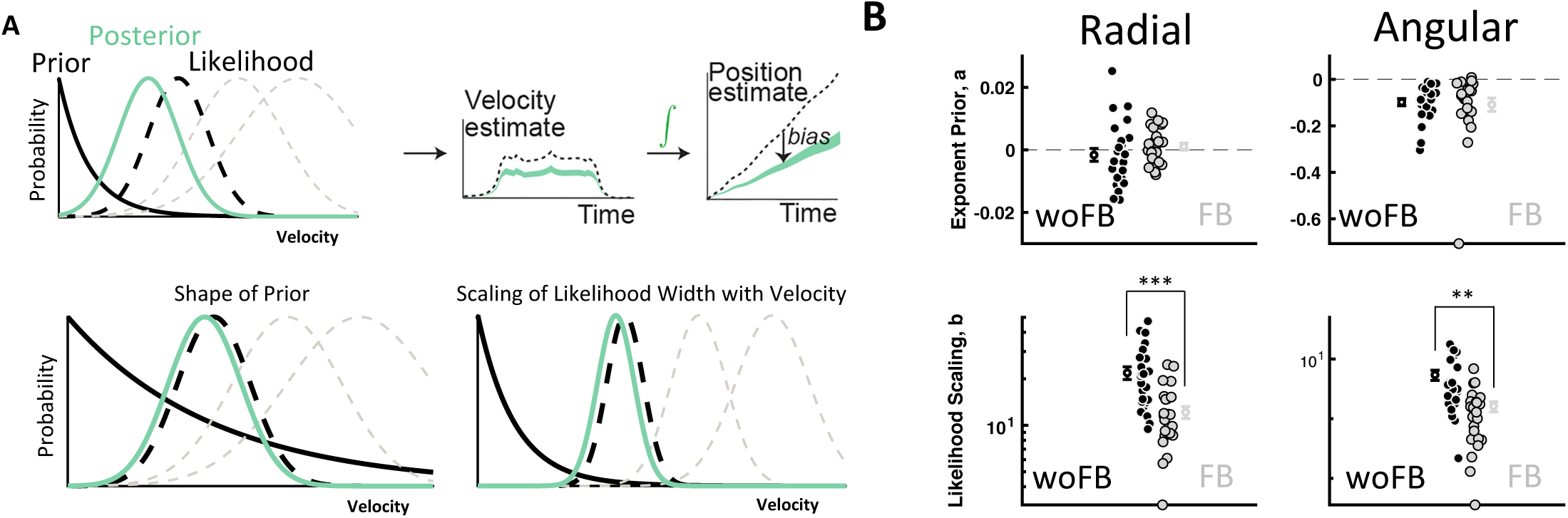
Speed-Prior Dynamic Bayesian Observer Model of Path Integration. **A)** Biases in path integration originate from an underestimation of velocity, modeled as a posterior (in green) based on a prior for speeds (black solid line) and a likelihood distribution (black dashed line). The likelihood width is taken to scale with velocity, as shown by the superimposed gray likelihoods increasing with width. This velocity posterior is then integrated into an estimate of position. Improvement in performance may be due to either the prior relaxing (bottom left), or the scaling of likelihood width becoming shallower. **B)** Extraction of the parameters best accounting for participant trajectories suggests that the prior does not change with feedback, but instead the scaling of the likelihood width with velocity becomes shallower.

The second, ‘leaky integration’ model (**Figure S2B**), hypothesizes that estimates of velocity are correct, and instead biases are due to imperfect integration of velocity into position estimates (Lappe et al., 2007, 2011; Mittelstaed & Glasauer, 1991). This assumes a flat prior (*a*_*v*_ = *a*_*w*_ = 0), but a leaky integration. Integration is dictated by two independent leak time constants *τ*_*d*_ and *τ*_*φ*_ that specify the timescale over which radial and angular velocity are integrated. This ‘leaky integration’ model was refuted for neurotypical individuals by Lakshminarasimhan et al., 2018, but has been included here to compare between ASD and controls, given that one of the many ASD hypotheses claims deficient integration in this conditions (Iarocci & McDonald, 2006; Stevenson et al., 2014; Wallace et al., 2019).

Both the speed prior and leaky integrator models have 4 free parameters; parameters *b*_*v*_ and *b*_*w*_ expressing how fast the squared spread (i.e., variance) of the likelihood functions scale with the magnitude of linear and angular velocity measurements, and either *a*_*v*_ and *a*_*w*_ (for the speed prior model) or *τ*_*d*_ and *τ*_*φ*_ (for the leaky integrator model). To gauge the quality of model fits, we reasoned that a good behavioral model ought to believe that the participants *should* stop where they *did* stop. In other words, the model belief should demonstrate that participants stopped where they did because they believed to be near the target. In turn, if the model accounts well for participant’s beliefs, its position estimate should be concentrated near the true target. To evaluate this, we used the best-fit model parameters for each subject to reconstruct the subjects believed end position. There was no residual bias (i.e., bias after accounting for the subject’s best fit parameters), neither prior to nor after feedback, and neither in the radial nor angular dimension, when utilizing the speed-prior model (“residual” bias contrast to a slope of 1 – no bias – all p > 0.10; **Figure S2A**). On the other hand, under the architecture of leaky integration, all contrasts showed significant residual biases (all p < 0.02; **Figure S2B**).

Given that the speed prior model (**Figure 2A**) accounted best for the observed data (both in the conditions with and without feedback), there are two putative mechanisms leading to enhanced performance in the block with feedback. The first is that during the block with feedback, the exponential prior for speed relaxes, becomes closer to a uniform prior (**Figure 2A**, bottom left). The second is that the scaling of the variance of the likelihood distribution with velocity becomes shallower (**Figure 2A**, bottom right). To distinguish between these possibilities we extracted the latent parameters of the best-fit model for each participant, which were then compared across feedback conditions. As shown in **Figure 2B** (top), there was no change with feedback in the exponent characterizing the prior for speed, neither in the radial (ΔRadial = 0.002 ± 0.002, p = 0.2) nor angular (ΔAngular = - 0.01 ± 0.014, p = 0.71) dimension. Of note, however, while the angular exponent was on average (i.e., taking into account both the block with and without feedback) significantly smaller than zero (- 0.10 ± 0.02, p = 1.0 × 10^−7^), the radial one was not (−2.75 × 10^−4^ ± 0.001, p = 0.81), suggesting that the greater bias exhibited by participants in the angular than radial dimension in the block without feedback is driven by a stronger prior in the former dimension.

On the other hand, the scaling of the likelihood variance with velocity decreased with feedback, both in the radial (ΔRadial = −9.69 ± 2.0, p = 9.9 × 10^−5^) and angular (ΔAngular = −3.81 ± 0.96 p = 7.30 × 10^−4^) dimension (**Figure 2B**, bottom). Together, the modeling results indicate that biases in path integration originate chiefly from biases in velocity estimates that are then appropriately integrated into position estimates. On the other hand, the improvement in path integration accuracy is driven by a reduction in the scaling of uncertainty with velocity, and not in a change in the prior.

### Abnormal Uncertainty Prior to Feedback in Autism

Next, we assess path integration abilities in individuals with ASD, and examine whether feedback improves their performance using a strategy akin to that employed by neurotypical individuals. We start with a model-independent quantification of the behavior, and end with a direct comparison with Bayesian model fits.

Prior to feedback, the clinical group showed marked overshooting of targets, both in the radial (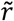 vs. *r* slope = 1.10 ± 0.08, p = 0.048) and angular (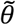 vs. *θ* slope = 1.52 ± 0.07, p = 1.3 × 10^−5^) dimension. The magnitude of this bias was not different than that of neurotypical controls (all p > 0.73, **Figure S3**, y-axis). Providing feedback at the conclusion of each trajectory improved their performance (feedback – without feedback, ΔRadial = −0.12 ± 0.08, p = 0.021; ΔAngular = −0.40 ± 0.07, p = 2.2 × 10^−4^), and the magnitude of this enhancement was similar across control and ASD groups (all p > 0.92; **Figure S3**, x-axis). Similar to the control subjects, during feedback individuals with ASD shortened their trial durations (p = 7.69 × 10^−9^) by increasing radial (p = 2.71 × 10^7^) and angular velocities (p = 0.01, **Figure S1**). In fact, they employed this same strategy to a greater extent than their neurotypical counterparts (all p<0.045). Further, the trial-to-trial dynamics with which feedback improved end-point accuracy within these groups was ostensibly also similar, as suggested by the fact that the outer boundary of the “rewarded zone” decreased at the same rate (see **Figure S4**).

Contrary to the similarity in performance when average responses were considered, indexing of dispersion tendencies suggested that at baseline (i.e., prior to feedback) individuals with ASD exhibited a heightened degree of variability with respect to control individuals. Namely, prior to feedback, trial-to-trial variance in radial endpoints, as captured by R^2^-values of the linear regression fits, was larger in ASD than control individuals (p = 1.3 × 10^−6^; **Figure 3A**). The introduction of feedback eliminated the deficit shown by the clinical group (p = 0.61). A similar analysis contrasting the variability of angular responses during the no feedback block did not detect a significant difference between groups (p = 0.36, **Figure 3B**), possibly due to a ceiling effect (77 / 78 R^2^-values > 0.92). To circumvent this problem, we also quantify uncertainty as the standard deviation of a select group of trials, split into equally sized quartiles based on target distance. This analysis showed that prior to feedback endpoint responses of ASD individuals were more variable than that of control participants (**Figure 3C, D**) both in the radial (p = 0.030) and angular (p = 0.048) dimension. Further, uncertainty scaled with target distance (both p < 0.001), a scaling that was exacerbated in ASD for the radial dimension (p = 0.028). Feedback reduced variability (both p < 0.001), equating the clinical and control group (**Figure 3C, D**). Importantly, the correlation between target distance and increasing uncertainty – and the greater scaling of the latter with the former in the ASD group – was true when conducting partial correlations accounting for response distance, trial duration, movement duration, reaction time, and mean movement velocity (p < 0.05). Thus, the greater variance in ASD is not merely motor, where response duration or response distance would have accounted for the variance instead accounted by target distance.

**Figure 3.**
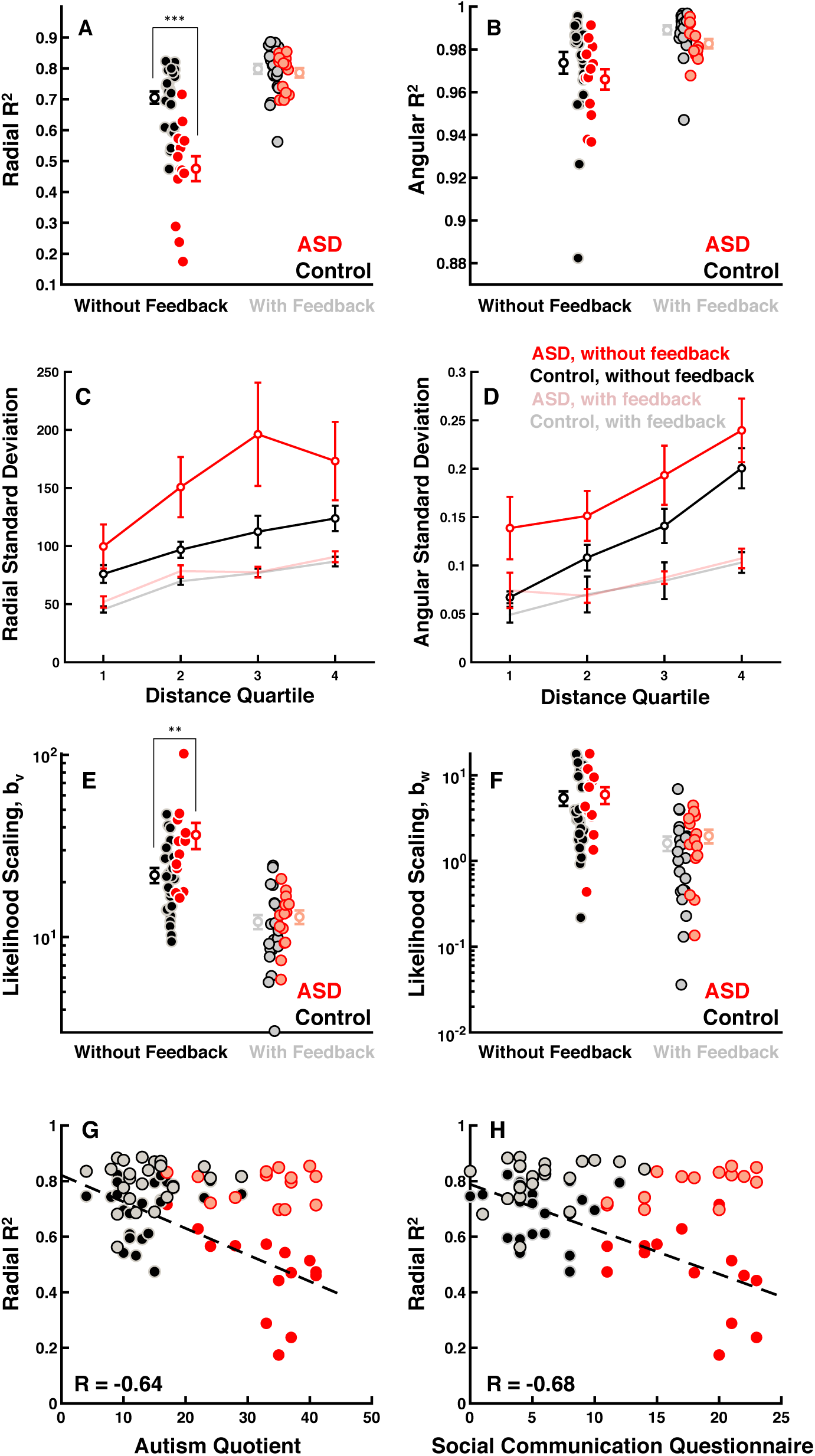
Increased Uncertainty in Autism. **A, B)** Goodness-of-fit (R^2^) of linear regression between response vs. target distance (from plots as in Fig. 1E-H) for ASD (red) and control (black) subjects without and with feedback. Data shown for individual subjects and group averages (+/-S.E.M.). **C**,**D)** Standard deviation of the end-point responses within specific target distance bins (x-axis) in the radial (left) and angular (right) dimension. **E, F**) Variance of the likelihood function (computed from the speed-prior model fit). **G**,**H)**, Radial R^2^ correlate inversely with ASD symptomatology prior to feedback (solid colors and dashed lines): the larger the endpoint variability, the higher participants scored on the Autism Quotient and the Social Communication Questionnaire. Single dots are individual participants.

As in the case with the control individuals, we leveraged the full extent of movement trajectories in 2-dimensions to track belief states by fitting the two abovementioned dynamic Bayesian Observer Models; one hypothesizing a prior for speeds leading to inaccurate velocity estimates, and the other hypothesizing an imperfect process of integration. As for the control subjects, the model that accounted best for participants’ trajectories was the speed prior (**Figure S2**). These results refute that ASD individuals are poorer than controls in integrating visual velocity signals over time (see Zaidel et al., 2015 for a similar conclusion using a passive visual motion integration task). Importantly, the best-fit estimated speed prior was equal in ASD and control groups both before (Linear, p = 0.61; Angular, p = 0.13) and after feedback (Linear, p = 0.42; Angular, p = 0.89; see **Figure S5**). In contrast, and as expected from the R^2^ results, the best-fit parameters of the scaling of the radial likelihood variance with velocity reflected a steeper dependence (i.e., uncertainty grew faster with velocity) in ASD than controls (p = 0.01, **Figure 3E**). The difference in radial likelihood scaling between ASD and controls disappeared after feedback (p = 0.64). While generally the scaling of the angular likelihood with velocity also decreased with feedback in the ASD group (p = 2.93 × 10^−5^), it did not do so differently than for neurotypical controls (all p > 0.67, **Figure 3F**).

What is perhaps most notable is that the degree of trial-to-trial variability in radial endpoints before feedback was not only different across experimental groups, but also showed strong correlations with ASD symptomatology. The Autism Quotient (AQ; Baron-Cohen, 2001), measuring symptoms of autism spectrum in healthy populations, and the Social Communication Questionnaire (SCQ; Rutter et al., 2003), measuring communication skills and social functioning symptomatology, both negatively correlated with the R^2^-values of the linear regression between radial targets and responses prior to feedback (radial R^2^ and AQ, r = - 0.64, p = 9.52 × 10^−6^; radial R^2^ and SCQ, r = - 0.68, p = 1.5 × 10^−6^; **Figure 3G, H**). Further supporting the association between greater variability and ASD symptomatology, the scaling of radial uncertainty with velocity extracted from the best-fit speed prior model also showed a positive correlation with both the AQ (r = 0.53, p = 4.63 × 10^−4^) and SCQ (r = 0.55, p = 2.27 × 10^−4^; **Figure S6)**. These findings consistently suggest that worsened symptomatology is associated with heightened variability in responses.

The observed differences in uncertainty between ASD and control groups, as well as the association of this variability with ASD symptomatology also extended into finer grain inspection of the trajectory themselves. Namely, as expected given the fact that trials begin and end with null velocities, radial distance from origin as a function of time was very well described by sigmoidal functions (r = 0.99 ± 6.34 × 10^−4^; **Figure 4A**; see Methods). These trajectories, however, were smoother in control than ASD individuals (p=0.001; **Figure 4A** shows a handful of example trajectories in a control and ASD participant, blue arrows indicate moment of jerkiness). Notably, unlike end-point variability, the quality of this fit did not change with feedback for either group (p = 0.13; **Figure 4B**). But importantly, the smoothness of movement trajectories in virtual reality scaled with ASD symptomatology: higher AQ (r = −0.42, p = 0.007, **Figure 4C**) and SCQ (r = −.50, p = 8.63 × 10^−4^, **Figure 4D**) scores were associated with larger deviations from perfectly sigmoidal trajectories both prior to and after feedback.

**Figure 4.**
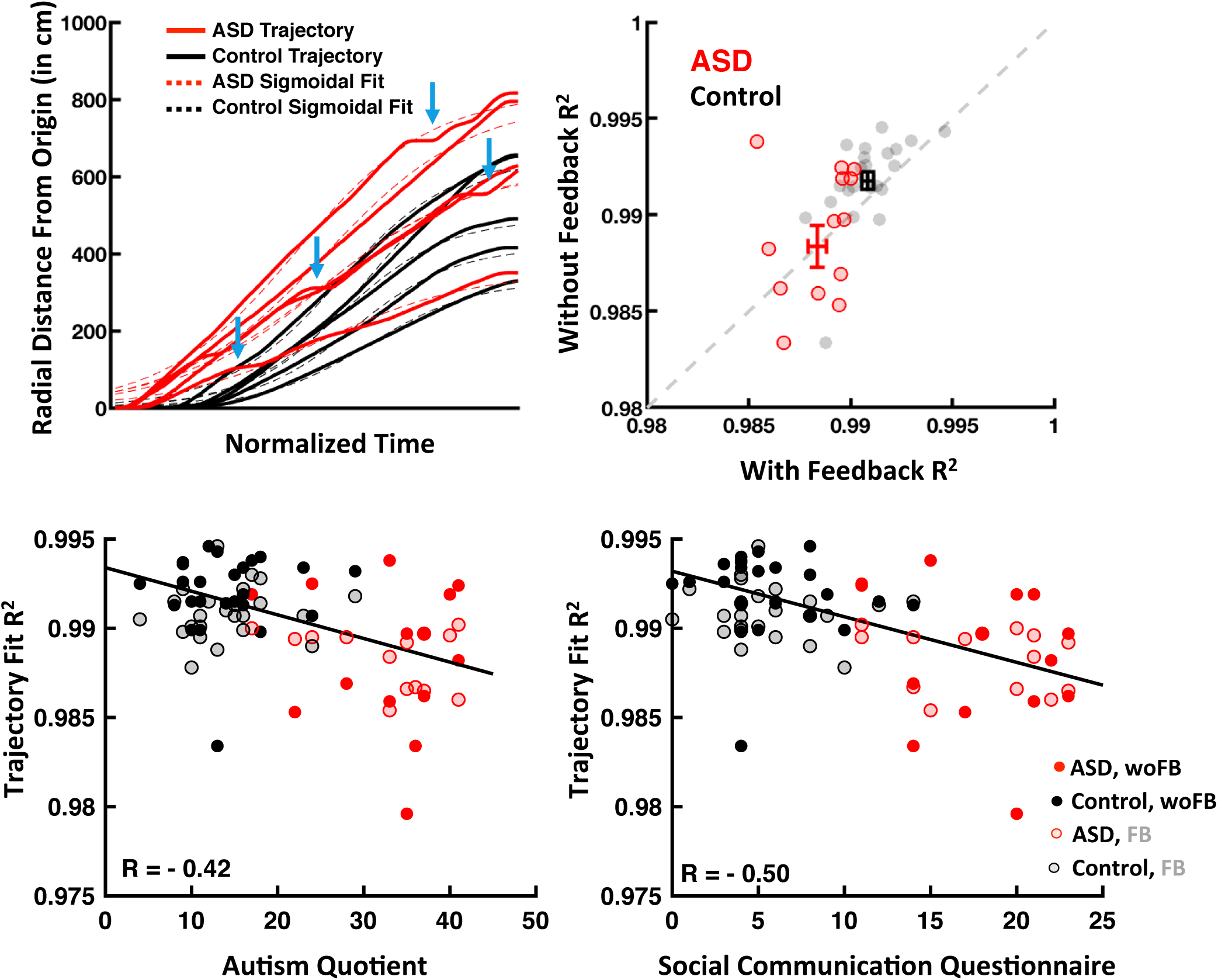
Trajectory smoothness correlates inversely with ASD symptomatology. **A)** Radial distance from the origin as a function of time (solid lines) was fit with a sigmoidal function (dashed lines); shown for a handful of example trajectories from a control (black) and ASD (red) individual. Blue arrows mark examples of jerkiness in ASD trajectories. **B)** Scatter plot of R^2^ of the sigmoidal fit with and without feedback for ASD (red) and control (black) individuals (also shown are means +/- S.E.M). **C**,**D)** The R^2^-values of sigmoidal fits correlated with both the Autism Quotient (AQ) and Social Communication Questionnaire (SCQ), suggesting that the smoother a participant’s trajectory, the lower they scored on ASD-related symptomatology..

## Discussion

Using a dynamic action-perception integration task, we found that on average ASD and control populations performed similarly, showing comparable biases (target overshooting; Lakshminarasimhan et al., 2018) in the absence of feedback. Similarly, both groups were also able to ameliorate their performance given feedback at the end of each trial. The initial bias seemingly stemmed from a speed prior (Stocker & Simoncelli, 2006; Petzschner & Glasauer, 2011; Lakshminarasimhan et al., 2018) biasing estimates of self-velocity, and not from the leaky integration of velocity into position estimates (Mittelstaedt & Glasauer, 1991; Lappe et al., 2007, 2011). Thus, the first conclusion of this work is that, contrary to what has been suggested in some previous literature (e.g., Noel et al., 2018; Iarocci & McDonald, 2006), individuals with ASD do not appear to be particularly poor integrators, at least not within the cadre of a naturalistic task wherein the integration is across a sustained time-period on the order of 5-6 seconds. This finding is in line with Giovanni et al., 2009, who demonstrated no difference between children with ASD and control individuals in blindly navigating toward a briefly presented target, using nothing but proprioceptive and kinematic information (as opposed to visual flow here). This finding is also in line with Zaidel et al., 2015, who demonstrated that both visual motion integration and the (maximum likelihood estimation, Ernst & Banks, 2002) integration of visual and vestibular signals into a multisensory estimate of self-motion was optimal and no different across control and ASD individuals. Overall, the current findings question theories of ASD emphasizing marked deficits in information integration (Happe & Frith, 1996, 2006; Iarocci & McDonald, 2006).

By fitting a dynamic Bayesian observer model across the entire trajectory, we were able to track participant’s belief states (Lee et al., 2014) and estimate likelihood functions and priors. The second conclusion of this work is that, contrary to the hypo-prior hypothesis of ASD (Pellicano & Burr, 2012), results showed no difference in the prior for self-motion speeds, neither in the radial nor angular dimension, and neither prior to or after feedback. Thus, the only two studies that have computationally disentangled individual subject’s likelihoods and priors found no difference in prior distributions between ASD and control groups (Karvelis et al., 2018; present study). These results contradict the hypothesis that hypo-priors can indeed represent a global property of ASD. Importantly, these results do not argue against the hierarchical Bayesian framework for ASD, rather they simply demonstrate that a canonical signature of ASD cannot be found in the rather superficial and simplistic hypo-prior explanation.

On the other hand, results revealed heightened variability in the clinical population. Namely, whether uncertainty was quantified as the fitted width of likelihood functions in the Bayesian model, by summary statistics (i.e., R^2^), or by examining the finer-grain detail of trajectories, we found that individuals with ASD were more variable than their neurotypical counterparts. Further, we found that across multiple task-variables (i.e., target distance and movement velocity) variability scaled quicker in ASD than in controls. Importantly, the variability of path integration endpoints (quantified by either R^2^ or likelihood widths within a dynamic Bayesian Observer model), as well as the degree to which movement trajectories deviated from smooth sigmoids, were all associated with increased severity in ASD symptomatology, as measured both by the AQ and SCQ scores.

Given the closed-loop nature of our task, this heightened variability can reflect either an impairment in filtering out noise on the sensory side or a decreased ability to generate consistent motor responses. The former explanation is in line with findings with simpler, open-loop tasks (Dinstein et al., 2012, Haigh et al., 2014; Bonneh et al., 2011; Milne, 2011; Zaidel et al. 2015). From a mechanistic standpoint, these widespread deficits in precision could emanate from a deficit in a global, brain-wide, computation. One of such canonical computations is divisive normalization (Carandini & Hegger, 2012), where neural firing is contextualized by a normalizing pool, effectively leading to a filtering operation. In fact, using neural network simulations, Rosenberg and colleagues (2015) demonstrated that anomalies in divisive normalization could account for an array of visual perception consequences reported in ASD, and more recently Coen-Cagli and colleagues (2019) showed that neurons that are more strongly normalized fire more reliably.

An alternative, but not necessarily mutually exclusive explanation, is that the increased variability in path trajectories and end points does not reflect a heightened sensitivity to noise at the stage of encoding, but rather an increase volatility of beliefs. Specifically, it has recently been proposed that the core abnormality in ASD may reside in perceptual aberrations due to an imbalance in predictive coding (Haker et al., 2016; Van de Cruys et al., 2016). According to this hypothesis, individuals with ASD overestimate their sensory prediction errors, such that the world appears more volatile than it is. This expansion of the original hypo-prior hypothesis to predictive coding (Friston et al., 2013) is not only qualitative consistent with previous experimental findings but has also been recently supported experimentally by Lawson and colleagues (2017), who showed that adults with ASD over-estimate volatility in the face of environmental change. This comes at the expense of learning to build stable expectations that lead to adaptive surprise, and thus leads to over-reacting to environmental change and being disproportionately receptive to sensory input. According to this framework, wrongfully precise prediction errors during inference would urge new learning that results in ‘noise’ as it is unlikely to repeat. Given the naturalistic, closed-loop action/perception/prediction nature of our task, where actions (driven by prediction errors) influence sensory inflow, such erroneous learning due to overestimated sensory prediction errors would lead to increased variability in actions, as observed here. With feedback, ASD performance matched that of control individuals, likely because feedback provided a veridical precision for sensory prediction errors. Thus, the present findings, which do not support the hypo-prior hypothesis, are qualitatively consistent with the sensory prediction error hypothesis.

While normative computational frameworks are certainly well positioned to account for the panoply of cognitive and perceptual abnormalities present in ASD, the recently promoted Bayesian approaches (Lawson et al., 2014, 2017; Palmer et al., 2017; Karvelis et al., 2018) are only a narrow view of a larger and more complex story. Notably, whether priors are weak (Pellicano & Burr, 2012) or too volatile (Lawson et al., 2014; Lieder et al., 2019), these theories emphasize anomalies that act exclusively at the decoding level – but this is but a small component of brain computation. To our knowledge, other aspects of statistical inference have so far been ignored; i.e., how likelihood functions are constrained by priors (i.e., efficient coding; Wei & Stocker, 2015). Our closed-loop task involves both encoding and decoding in an intertwined, naturalistic way, which may adhere better to ASD symptomatology of everyday experiences. We argue for the use of more naturalistic, dynamic tasks (e.g., where sensory processing, perceptual inference, and actions/decisions are not artificially segregated in laboratory tasks) and normative modeling to understand how the neural computation in individuals with ASD has gone awry.

Collectively, both studies that have explicitly fitted priors and likelihoods to participants’ behavior (Karvelis et al., 2018; present study) have demonstrated that the source of the computation that has gone awry in ASD is not the prior. Instead, a broader aberrant learning and inference may be the core component of maladaptive cognition within the condition. Maladaptive inference in ASD can arise from alterations in one of several core components (beyond priors and likelihoods of an extremely simplified Bayesian model) that span a multi-dimensional computational space (Haker et al., 2016). The hypothesized fundamental mechanism affected in ASD, hierarchical Bayesian learning, is much broader than just the simple equation argued in recent publications (Posterior = Prior x Likelihood). Sensory encoding, efficient coding (where the prior is an inherent component of sensory encoding; Wei & Stocker, 2015), action-oriented predictive coding (Clark, 2013; Friston 2009; Friston et al., 2010), all constitute critical components of cognitive computations that must be quantitatively explored in ASD. Although not exhaustive, the space of possibilities for how the phenotypic and clinical variability of ASD patients could arise from different impairments of hierarchical Bayesian inference is multidimensional. In other words, different autistic phenotypes could arise from different impairments in the computation of multiple key variables, which may qualitatively (but erroneously) appear as priors and likelihoods. A few years ago, a computational perspective on Autism was rare. Now, it is growing in appeal. However, we should not underestimate the complexity of both the brain and the ASD phenotype, particularly in light of the highly constrained experimental tasks and models we use.

## Methods

### Participants

Thirty-nine subjects completed the firefly catching task. Fourteen were individuals diagnosed within the Autism Spectrum Disorder (ASD; N = 14, mean ± sd; age = 14.5 ± 2.1 years; AQ = 32.7 ± 7.3; SCQ = 17.8 ± 4.2) by expert clinicians. The rest were age-matched neurotypical individuals (Control; N = 25, mean ± sd; age = 14.8 ± 2.14 years; AQ = 14.0 ± 5.5; SCQ = 5.6 ± 3.2). Participants had normal or corrected-to-normal vision, and no history of musculoskeletal or neurological disorders. Prior to partaking in the study, all participants completed the Autism Spectrum Quotient (AQ; Baron-Cohen, 2001) and the Social Communication Questionnaire (SCQ; Rutter al., 2003). The Institutional Review Board at Baylor College and Medicine approved this study, and all participants gave their written informed consent and/or assent.

### Materials and Procedures

Participants were tasked with virtually navigating to the location of a briefly presented target (i.e., the ‘firefly’) via an analog joystick with two degrees of freedom (linear and angular speed). Participants were seated facing a large projection screen (width × height: 149 × 127 cm) positioned 67.5cm in depth with respect to their eyes, and wore a seatbelt in order to restrain trunk movements. Visual stimuli were rendered as a red-green anaglyph and subjects wore goggles fitted with Kodak Wratten filters (red #29 and green #61) to view the stimulus. The virtual world comprised a ground plane whose textural elements were isosceles triangles (base x height = 8.5 × 18.5cm) that were randomly positioned and re-oriented at the end of their lifetime (lifetime = 250ms; floor density = 2.5 elements/m^2^; **Figure 1A**). This floor texture had a limited lifetime and was re-oriented at each presentation in order to allow them to provide optic flow information, but not serve as landmarks. The ground plane was circular with a radius of 70 meters (near and far clipping planes at 5cm and 4000cm, respectively), and the subject was positioned at the center of this virtual world at the beginning of each trial. On each trial the target was a circle of radius 20cm whose luminance was matched to the ground texture elements, blinked at 5Hz and appeared at a random location between θ = ± 42.5° of visual angle and at a distance of r = 1 – 6m relative to where the subject was stationed at the beginning of the trial. After one second, the target disappeared, which cued subjects that they could use the joystick to navigate to the location of the target. The maximum linear and angular speeds were respectively limited to *v*_*max*_ = 2 *m/s* and *w*_*max*_ = 90°/*s*. Upon arriving at the location where they thought the firefly was present, participants pressed a button to indicate their response.

The experiment consisted of 2 blocks, each block consisting of 150 trials. In the second block participants were given visual feedback. This feedback was in the form of a bull’s-eye pattern rendered on the virtual floor (**Figure 1B**). This pattern consisted of six concentric circles, with the radius of the outermost circle being continuously scaled (up or down by 5%) according to the 1-up-2-down staircase procedure. Additionally, an arrowhead indicating the target location was presented on the ground with either green or red color, depending on whether the participant’s final response was within or outside the outermost “rewarded” concentric circle (**Figure 1B**). The two blocks of trials (without and with feedback) were separated by at least 5 minutes rest.

All stimuli were generated and rendered using C++ Open Graphics Library (OpenGL) by continuously repositioning the camera based on joystick inputs to update the visual scene at 60 Hz. The camera was positioned at a height of 1 m above the ground plane. Spike2 software (Cambridge Electronic Design Ltd.) was used to record and store the subject’s linear and angular velocities 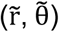, target locations (r, θ), and all event markers for offline analysis at a sampling rate of 833.3 Hz. Further details of the experimental setup and task can be found in Lakshminarasimhan et al. (2018).

### Data Analyses

The location of randomly presented targets (**Figure 1C**, left), participants’ trajectory (**Figure 1C**, right) and final position responses were expressed in polar coordinates as a radial distance (target = r; 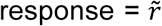) and an angular eccentricity (target = θ; 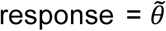; arbitrarily, straight ahead = 0°; **Figure 1C**). When visualizing responses as a function of target location (**Figure 1D**, right), it was apparent that a linear model with multiplicative gain scaling accounted well for the observed data (see *Results*, **Figure 1D**, error was greater at the farthest locations tested). Thus, we used the slopes of the corresponding linear regressions as a measure of bias. Note that in this schema a slope of 1 indicates no bias, while slopes larger than 1 indicate overshooting (either in radial distance or angle). For each subject we extract R^2^ values of the linear fit of radial/angular target vs. response as an indication of trial-to-trial variability. Lastly, virtual path trajectories were fit with a sigmoidal function (given that participants started and ended their trajectories at velocity v = 0; parameter dictating the location and steepness of the non-linearity were left as free parameters, the saturation points were taken to be the minimum and maximum observed in data). Trajectories were down-sampled to 83.33Hz prior to sigmoidal fitting.

### Dynamic Bayesian Observer Model

To account for the pattern of behavioral results, we considered an observer model comprised of a Bayesian estimator that uses noisy measurements *m*_*v*_ and *m*_*w*_ to decode linear and angular self-motion velocities *v* and *w*. These internal velocity beliefs were then temporally integrated to dynamically update the subject’s position in the virtual world. We parameterized the model by making the following three assumptions. First, we chose an exponential function to describe the priors over both linear and angular velocities: 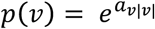 and 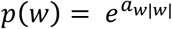. Second, likelihood functions *p*(*m*_*v*_|*v*) and *p*(*m*_*w*_|*w*) were assumed to be Gaussian, centered on the respective measurements *m*_*v*_ and *m*_*w*_. That is, likelihoods were unbiased. The variance of these likelihood functions scales proportional to the magnitude of velocity measurements: *Var*(*m*_*v*_)*= b*_*v*_ |*m*_*v*_| and *Var*(*m*_*w*_) = *b*_*w*_ |*m*_*w*_|. Under these conditions, it can be shown that the means and variances of the maximum a posteriori estimates 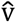 and 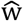 are given by (Stocker & Simoncelli, 2006; Lakshminarasimhan et al., 2018):

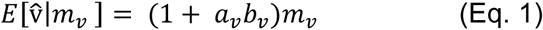

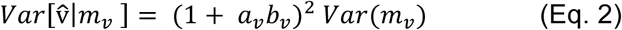

and correspondingly for 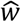. A flat prior corresponds to an exponent of zero yielding an unbiased estimate, while negative/positive values of the exponents would result in under/overestimation of the speeds.

The third and final building block of the model pertains to the integrator computing position from velocity. We assume that the integrative process is dictated by two independent leak time constants *τ*_*d*_ and *τ*_*φ*_ that specify the timescales of integration of estimated linear and angular speeds to compute distance (*d*) and heading (*φ*). In turn:

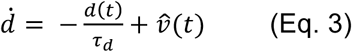

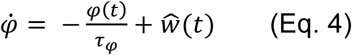

The average distance and heading at each time point can be determined by convolving the mean velocity estimates with an exponential kernel 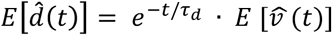 and 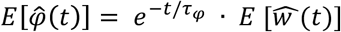, where the expectations are taken over the corresponding posterior probability distributions. Likewise, if the noise in the velocity measurements is temporally uncorrelated, the variance of the distance and heading estimates can be expressed in terms of the variances of the velocity estimates; 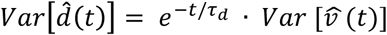 and 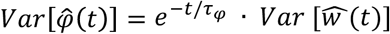. Hence, in this case, both mean and variance of the integrated estimates will share the same temporal dynamics. Note that the mean estimates 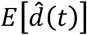 and 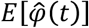 will be accurate with large time constants, but will be misestimated if these constants are comparable with travel time, *t*. Since position is determined jointly by the time course of distance and heading, it follows that the subject’s mean estimate of their linear and angular positions 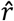 and 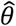 will also be different from their veridical values when *τ* ≈ *t.*

### Model Fitting

In a prior study our group (Lakshminarasimhan et al., 2018) demonstrated that a slow-speed prior model (i.e., with a negative *a*_*v*_ and *a*_*w*_ exponent), with perfect integration (*τ*_*d*_ and *τ*_*φ*_ set to infinity), best accounted for overshooting observed in this path integration task. Differently form Lakshminarasimhan et al., 2018, however, we allow for *a*_*v*_ and *a*_*w*_ to be either negative or positive (or zero; flat prior). In addition, as many (Rinehart et al., 2000; Iarocci et al., 2006; Robertson et al., 2012; Stevenson et al., 2014; Noel et al., 2018) claim that individuals with ASD are poor integrators, we also fit a leaky integrator model, where the prior was held flat (*a*_*v*_ = *a*_*w*_ = 0), yet *τ*_*d*_ and *τ*_*φ*_ were free parameters. Both the slow-speed prior model and the leaky integrator model have 4 free parameters; width parameters *b*_*v*_ and *b*_*w*_ expressing how fast the spread of the likelihood functions scale with the magnitude of linear and angular velocity measurements, and either *a*_*v*_ and *a*_*w*_ (for the speed prior model) or *τ*_*d*_ and *τ*_*φ*_ (for the leaky integrator model).

Since subject’s position estimates are probabilistic, we fit model parameters by taking both mean and uncertainty of position into account; this was done by maximizing the expected reward, that is, the probability that the subjects believed themselves to be within the target at the end of each trial. Model parameters were optimized utilizing MATLAB’s *fmincom* function, constraining time-constants (where applicable) and likelihood widths to be non-negative, and by initializing a total of 100 random seeds. For further detail see Lakshminarasimhan et al., 2018.

## Acknowledgements

We thank Jing Lin and Jian Chen for programming the experimental stimulus. This work was supported by the Simons Foundation, SFARI Grant 396921 and Grant 542949-SCGB, as well as R01 DC014678.

## Author contributions

DEA and KL designed experiments. HP collected data. JPN analyzed data. DEA and JPN wrote manuscript.

## Supplementary Materials

**Figure S1.**
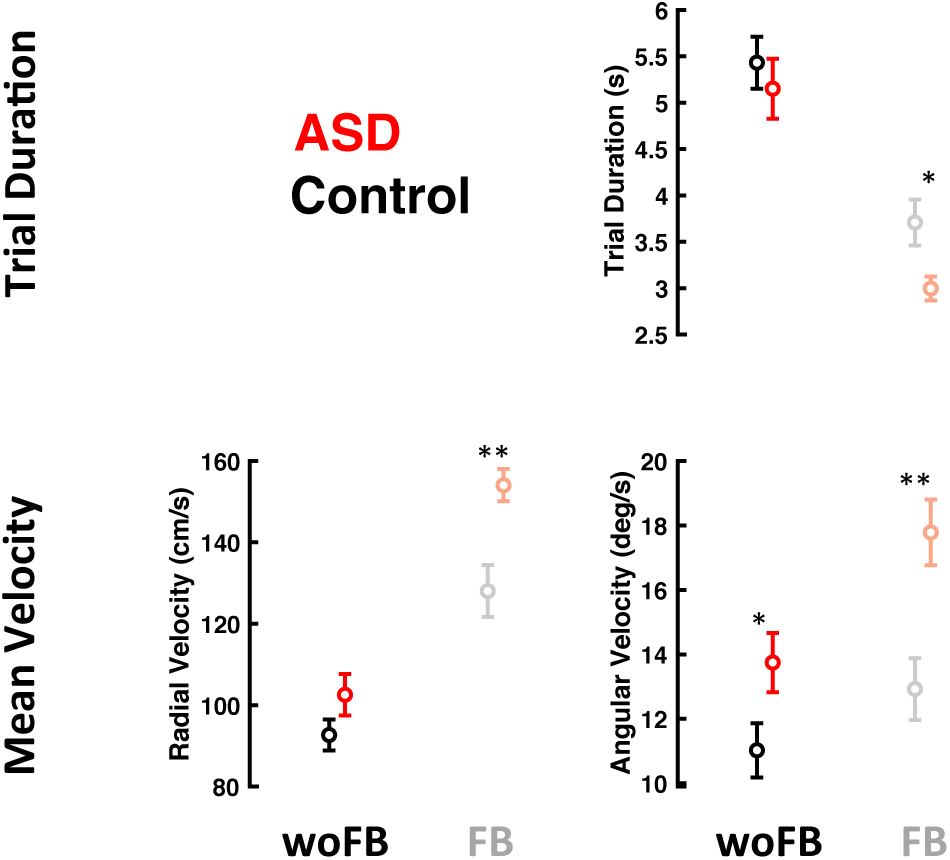
Trial duration decreases and mean velocity increases during feedback. Both ASD (red) and control (black) participants seemingly adopt a similar strategy in the blocks with feedback; they shorten their trials by increasing velocity. Statistics in main text, error bars are *± 1 SEM.*

**Figure S2.**
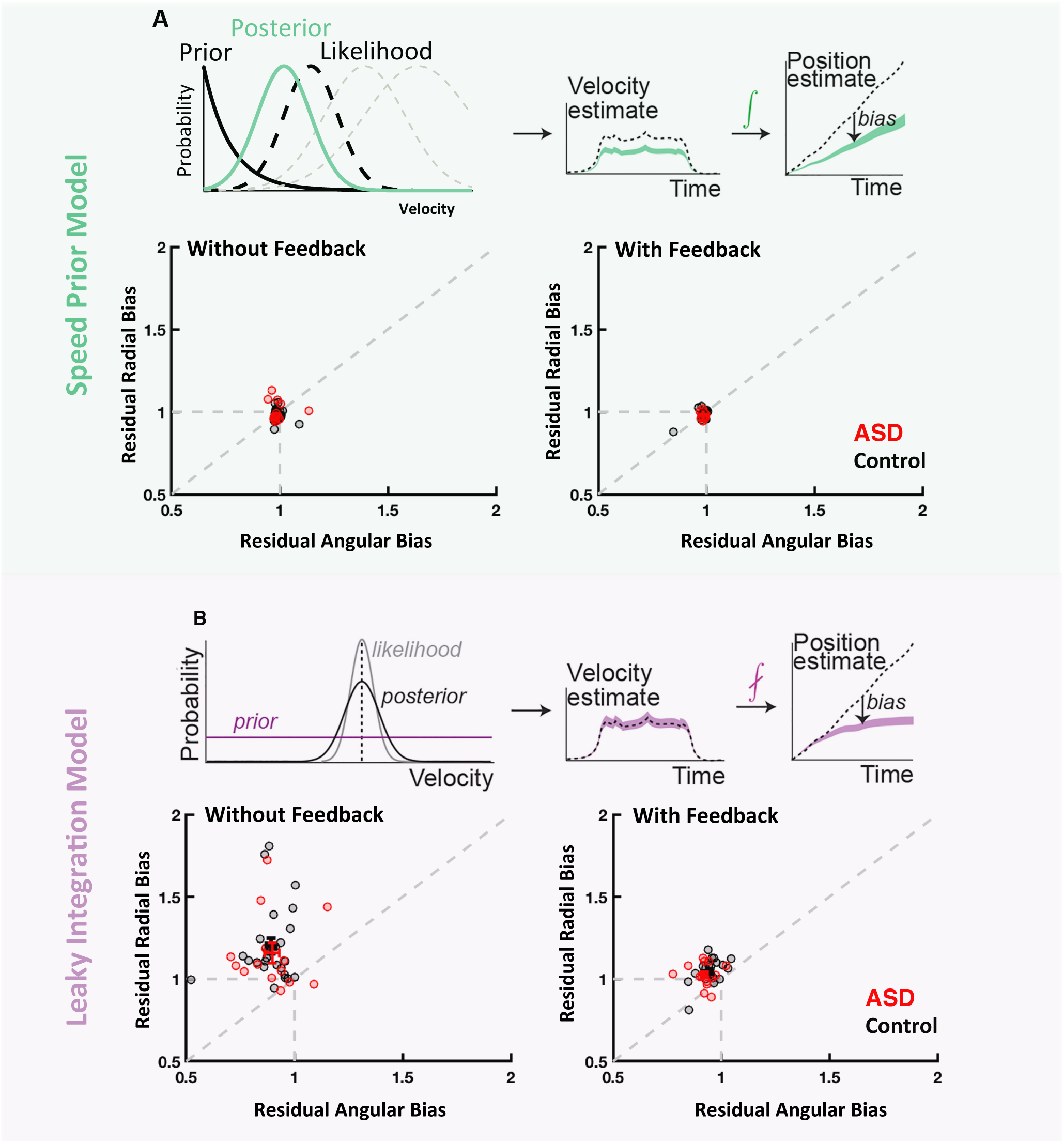
Dynamic Bayesian Observer Models. A) In the speed prior model, it is postulated that target overshoots are driven by a prior for speeds biasing velocity estimates. This prior (solid black) is combined with a sensory likelihood (dashed black and gray) to form a posterior estimate of velocity (green). This velocity estimate is estimated at each time-point (as likelihoods change) and integrated to yield an estimate of self-position. B) In the leaky integration model, it is assumed that velocity estimates are accurate (middle panel: purple line overlaps dashed line), but instead imperfectly integrated to form a position estimate. The correct model should account for subjects stopping where they did, believing they had reached the firefly. Fitting the former model and using extracted parameters to generate subjects’ believed trajectories shows residual biases (bias in model estimates) near 1 (i.e., no bias; A, lower panels). In contrast, the leaky integration model predicts large residual biases (B, lower panels), particularly failing to account for radial biases. Performance of the two models is comparable for ASD (red) and control (black) subjects, indicating that ASD subjects can perform sensory (path) integration as well as controls. Speed prior model: all residual biased, p > 0.15; leaky integration model: all biases significant except for angular bias with feedback, p < 0.02; No contrast in residual bias between experimental groups was significant (all p > 0.14)

**Figure S3.**
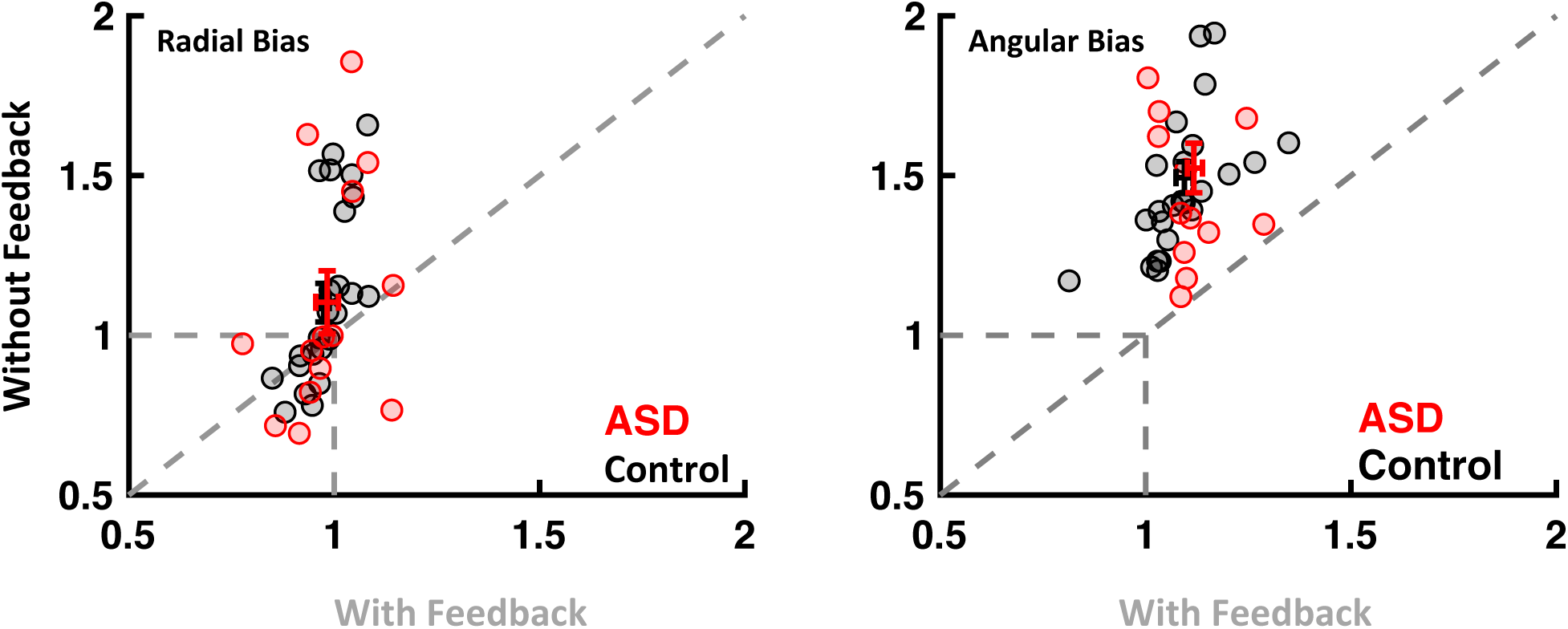
Effect of feedback in bias. Bias in path integration along the radial (left) and angular (right) dimension, without (y-axis) and with feedback (x-axis), and in control individuals (black) as well as individuals with ASD (red). As reported in the main text, both radial and angular dimensions showed significant overshooting (as shown by the average bias being above 1). Biases were reduced after feedback (as indicated by the averages being above the identity line). Importantly, prior to feedback radial (t-test ASD vs. control: p = 0.94) and angular (p = 0.62) biases were similar across the ASD and control groups. Similarly, the reduction in bias during the block with feedback was equal across groups, both for radial (t-test change in bias ASD vs. control: p = 0.945) and angular bias (p = 0.928). Error bars indicate *± 1 SEM.*

**Figure S4.**
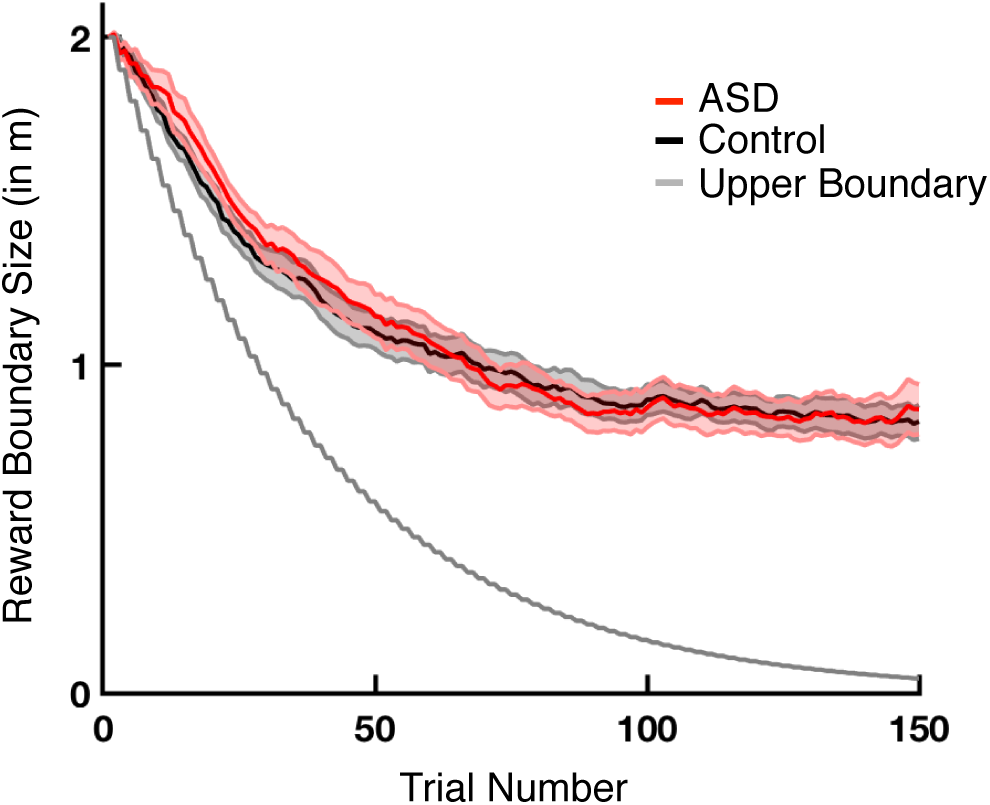
Trial-to-trial dynamics of the boundary demarking the rewarded zone during the feedback session. During the block with feedback, the “reward boundary size” decreased adaptively, starting at 2 meters. This boundary decreased with similar dynamics for both the ASD (red) and control (black) subjects. The decrease in the zone rewarded was far from the upper boundary of performance – the gray curve demonstrates the boundary of the rewarded zone if a participant were to fall within this zone on every trial (scales down by 5% upon two consecutive rewarded trials).

**Figure S5.**
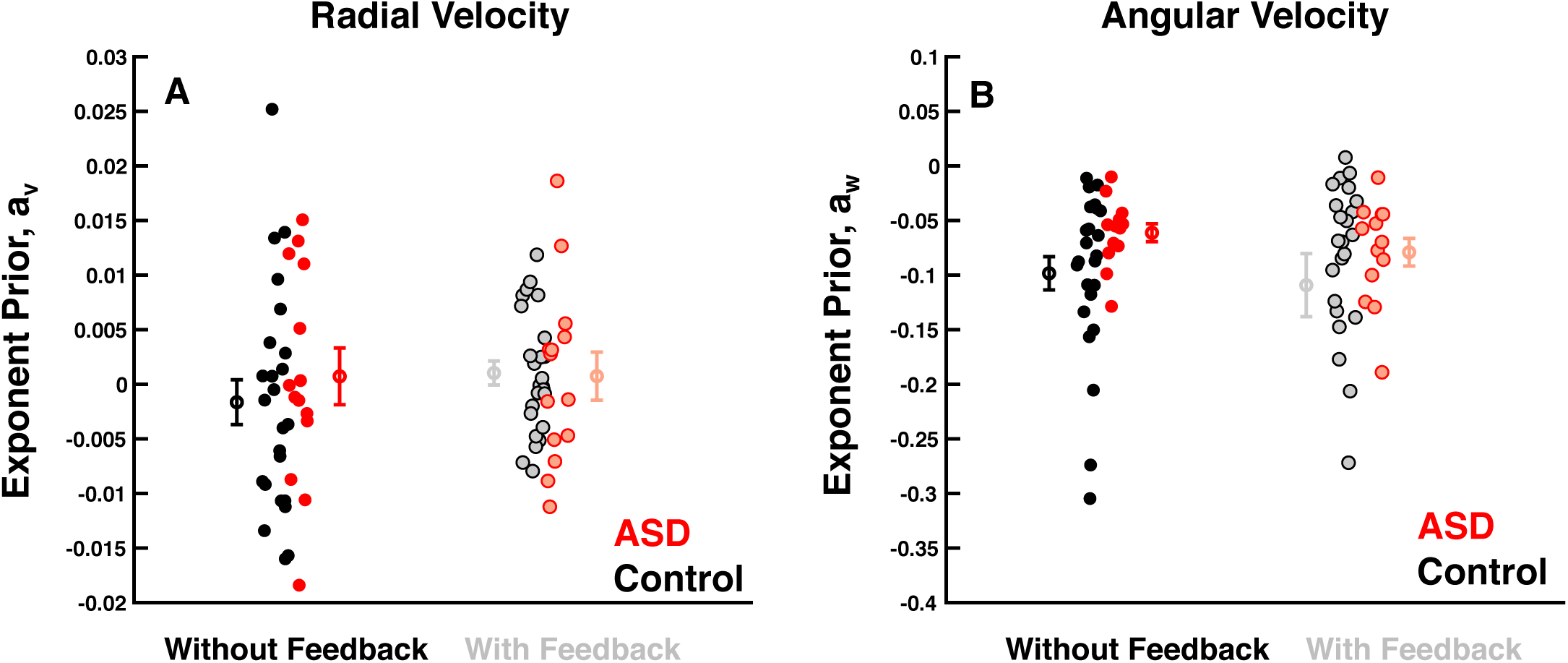
Speed Prior Parameter Estimates. The parameter dictating the shape of the exponential describing the prior over radial (**A**) and angular (**B**) velocity did not differ between experimental groups (ASD = red, control = black), nor feedback conditions. Exponential coefficients were not different from zero in the radial dimension, but negative (stronger overshooting biases) in the angular dimension.

**Figure S6.**
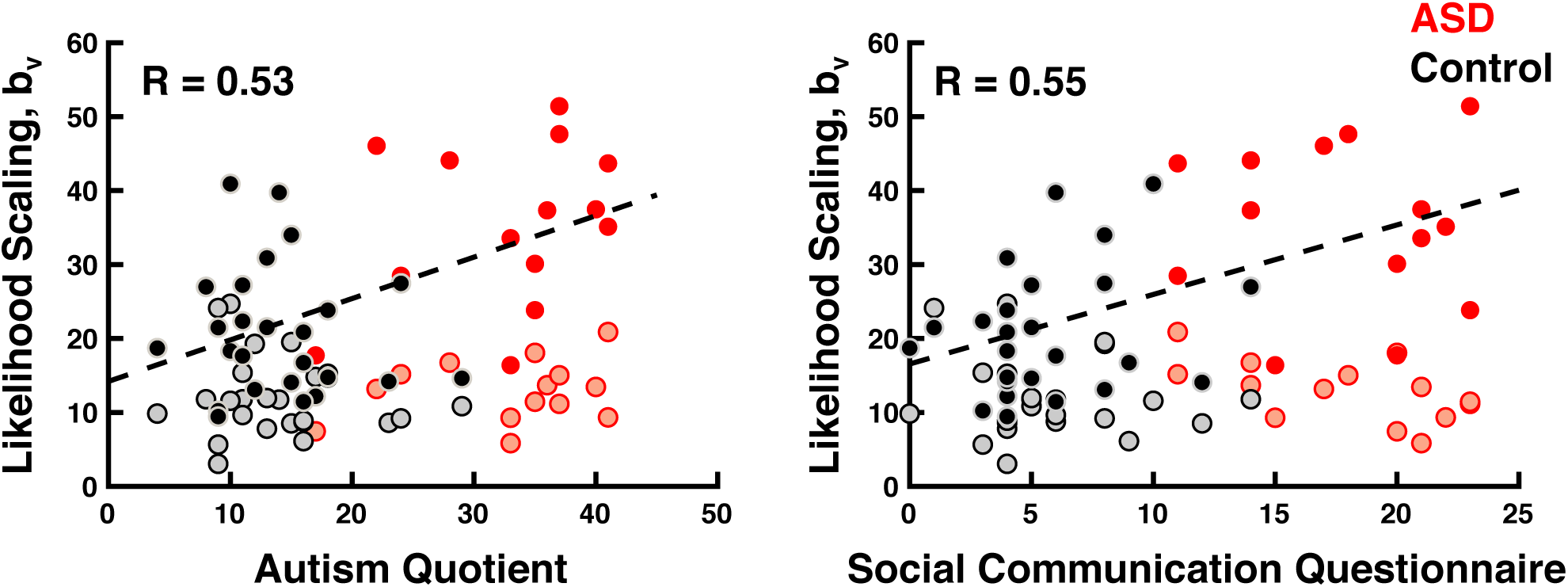
Likelihood scaling without feedback correlates with ASD symptomatology. Correlation between the estimated scaling of uncertainty with motion velocity according to the speed prior Bayesian dynamic observer model and ASD symptomatology. Both AQ (left) and SCQ (right) are higher for participants showing the greatest scaling of uncertainty with velocity in the block without feedback (solid filled symbols), but not after feedback (shaded symbols).

